# Predicting Individual Traits From T1-weighted Anatomical MRI Using the Xception CNN Architecture

**DOI:** 10.1101/2023.02.20.529226

**Authors:** Zvi Baratz, Yaniv Assaf

**Affiliations:** Department of Neurobiochemistry, Faculty of Life Sciences, Tel Aviv University, Israel

## Abstract

Modeling individual traits is a long-standing goal of neuroscientific research, as it allows us to gain a more profound understanding of the relationship between brain structure and individual variability. In this article, we used the Keras-Tuner library to evaluate the performance of a tuned Xception convolutional neural network (CNN) architecture in predicting sex and age from a sample of 4,049 T1-weighted anatomical MRI scans originating from 1,594 participants. In addition, we used the same tuning procedure to predict the big five inventory (BFI) personality traits for 415 participants (represented by 1,253 scans), and compared the results with those generated by applying transfer learning (TL) based on the models for sex and age. To minimize the effects of preprocessing procedures, scans were subjected exclusively to brain extraction and linear registration with the 2 mm MNI152 template. Our results suggest that CNNs trained with hyperparameter optimization could be used as an effective and accessible tool for predicting subject traits from anatomical MRI scans, and that TL shows potential for application across target domains. While BFI scores were not found not be predictable from T1-weighted scans, further research is required to assess other preprocessing and prediction workflows.

## Introduction

Computer vision has rapidly become an invaluable resource in both research and industry, with deep learning (DL) architectures, and particularly convolutional neural networks (CNNs), constituting the state-of-the-art (SOTA) for a wide range of classification and regression tasks. In many fields, such as object detection (Girshick, 2015), semantic segmentation (Badrinarayanan et al., 2017), and image denoising (Zhang et al., 2017), CNN architectures have been shown to deliver superior performance as opposed to any technique used in the past. New architectures are constantly emerging, with better and faster results, and DL models are quickly being integrated into industrial applications, effectively becoming the new standard for a growing number of commonplace tasks.

CNN models do not require manual feature engineering and can learn high-level abstractions of information in the data (Wu, 2017). However, in order for a model to be able to learn very fine details and precisely tune its weights to produce reliable predictions, many observations are required. So far, large data repositories have been mostly generated for 2D images, e.g., MNIST (for digit recognition) and ImageNet (for object classification). The aggregation of high-quality datasets at large scales is an arduous effort, demanding a massive investment in the manual curation and validation of every single observation. In addition to this technical challenge, medical data are subject to privacy constraints, which can serve to bar sharing and open distribution.

Despite these difficulties, a growing number of large MRI datasets has been made accessible (with varying degrees) to the research community. Notable examples include the UKBioBank (Bycroft et al., 2018), the Human Connectome Project (HCP) (Van Essen et al., 2012, 2013), the Autism Brain Imaging Data Exchange (ABIDE) (Di Martino et al., 2014, 2017), and more. As the scale of available data increases and deep neural networks (DNN) are becoming more readily trainable, they are also quickly finding their way into applications such as image segmentation (Akkus et al., 2017; Dalca et al., 2019) and registration (Andrade et al., 2018; X. Chen et al., 2021; de Vos et al., 2019; Shao et al., 2021; Xiao et al., 2021), tumor detection (Gore & Deshpande, 2020; Nazir et al., 2021; Siar & Teshnehlab, 2019), and even prediction of disease onset and progression (Liem et al., 2017; Liu et al., 2020; Lundervold & Lundervold, 2019; Noor et al., 2019; Saratxaga et al., 2021). CNN architectures may be built and optimized specifically for 3D neuroimaging data; however, there are numerous factors making a substantial difference in the estimated results. Such factors include differences in the acquisition parameters and environment, preprocessing tools and configurations, sample size, and the integrations of various data augmentation techniques.

To reduce the effect of preprocessing decisions on subsequent modeling efforts, it is enticing to use data that are as close as possible to their raw form and independently execute some chosen, uniform workflow. However, this approach is unscalable both in terms of the bandwidth required to acquire the data and the compute power necessary for other researchers to run preprocessing workflows with the computational resources which are generally available to conventional neuroimaging laboratories. For this reason, many public datasets make the results of some standardized preprocessing procedure available alongside, or even instead of, the raw data. In such cases, when combining results from different datasets, minor differences in workflows or configurations may have a dramatic effect on subsequent modeling efforts, and significantly hinder the reproducibility of derived work. As opposed to this, DNN architectures and weights may be applied to minimally preprocessed data, and any differences in acquisition or preprocessing configurations can be factored in. In addition, the encoded representations of individuals may be shared as simple vectors without any risk of privacy violations.

Even though large datasets are increasingly accessible, and building and training DNNs using high-level libraries is easier than ever before, there are still countless possible variations of architectures and training procedures. Choices such as the number of hidden layers, the number of filters, kernel sizes, regularization parameters, average versus maximum pooling, etc., all have a substantial effect on the performance of the network. Libraries such as Keras (see https://keras.io/) enable assigning and modifying all of these parameters with very few lines of code. This, again, introduces many degrees of freedom which hamper the ability of researchers to compare results across studies conducted on data from different scanning sites, using different acquisition protocols, on different populations, etc.

With Keras-Tuner’s HyperModel class (see https://keras.io/api/keras_tuner/hypermodels/), the hyperparameter search space can be declaratively integrated into the model building pipeline and facilitate a structured and reproducible framework for model training with minimal effort. In addition to its general utility in creating a robust and shareable workflow, the library also offers two prebuilt ResNet (He et al., 2016) and Xception (Chollet, 2017) hypermodel subclasses (HyperResNet and HyperXception, respectively). These provide well-tested and widely used implementations for these two types of CNN architectures, and enable easy adaptation and exploration in diverse subject domains.

An increasingly common method for using robust CNN architectures and taking advantage of previously trained weights in diverse contexts is through the application of Transfer Learning (TL). TL is a machine learning technique that involves using the knowledge gained from solving one problem to solve a related, but different problem (Torrey & Shavlik, 2009). In medical research, TL has been applied in several areas, including image classification (Godasu et al., 2020; Kora et al., 2022), natural language processing (Frei et al., 2022; Peng et al., 2019), and more (S. Chen et al., 2019). One of the main benefits of TL in medical research is that it allows researchers to leverage existing data and models, rather than starting from scratch, which can save time and resources. Additionally, TL can also improve the performance of machine learning models by transferring knowledge gained from one domain on to another, which can be especially useful in cases where there is a limited amount of data available for the target task.

In this article, Keras-Tuner’s built-in HyperXception CNN implementation was used to predict sex and age in the general population over a sample of 4,049 T1-weighted anatomical MRI scans from 1,594 subjects, as well as personality traits over a sample of 1,253 scans from 415 subjects. We then tested the success of TL applied between sex and age, trained on an identical sample, in comparison with the performance obtained when adapting the two models for the prediction of personality traits.

As researchers in an interdisciplinary field, leveraging relatively novel and highly complex technologies, it is imperative we continuously reassess the scalability and dependability of different ways to conduct our analyses and share our results. While the widespread adoption of DL techniques in neuroimaging is imminent as it is in many other fields, and is already showing promising results, it is important to emphasize and encourage the use of standardized, reviewed, and reusable methodologies. We hope the results of this study will benefit this larger effort.

## Materials and Methods

### Participants

This study uses a dataset collected from 1,594 participants (750 female and 844 male), 18 – 87 years old (mean: 31.02, SD: 10.82; see figure 1 for an overview of participant age distribution by sex). Participants were neurologically and radiologically healthy, with no history of neurological disease, and normal appearance in a clinical MRI protocol. The imaging protocol was approved by the Institutional Review Boards of Sheba Tel HaShomer Medical Center and Tel Aviv University, where the MRI investigations were performed. All participants signed informed consent forms.

**Figure 1.**
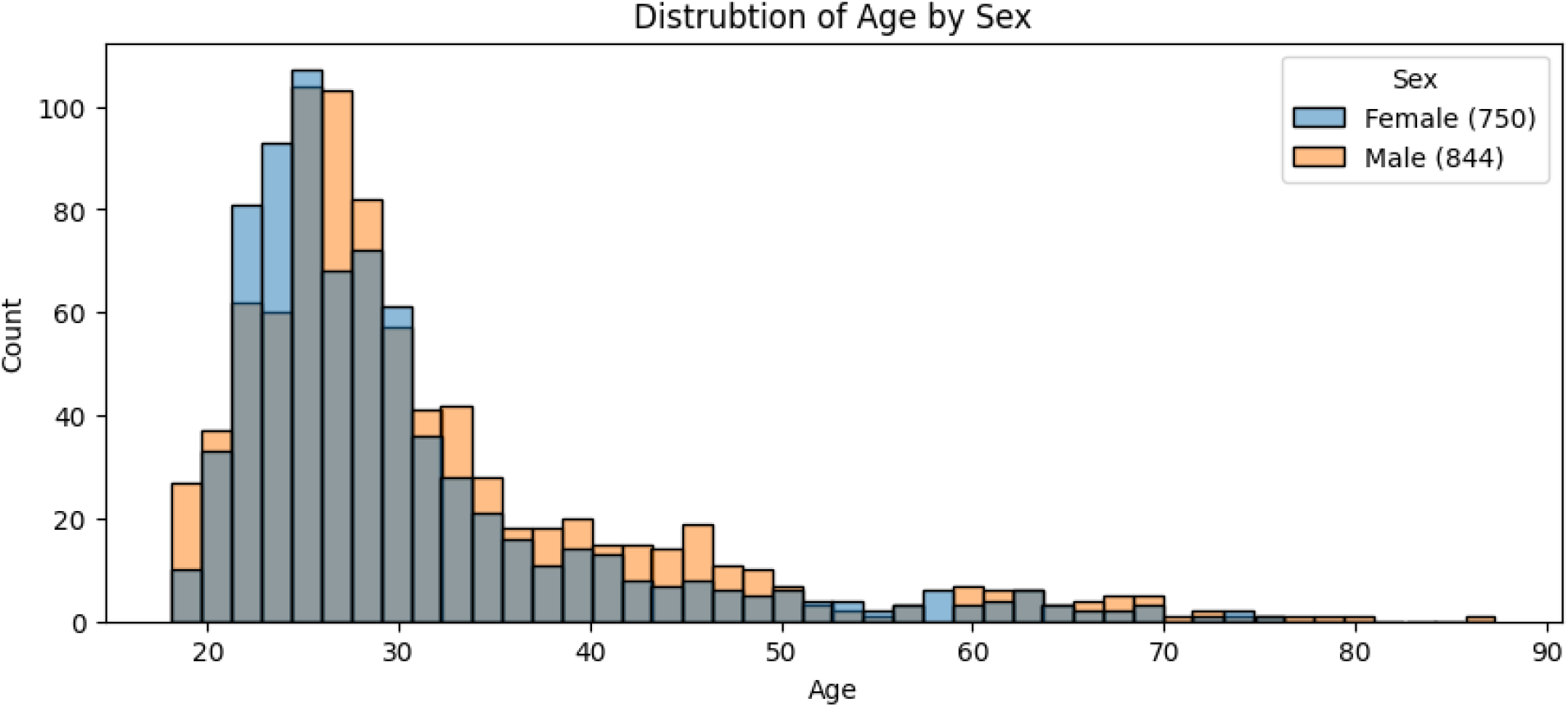
Distribution of participant age by sex. Total counts are denoted in parentheses within the legend.

### MRI Acquisition

All participants were scanned at the Alfredo Federico Strauss Center for Computational Neuroimaging at Tel Aviv University. Scans were acquired with a 3T Siemens MAGNETOM Prisma MRI scanner (Siemens Medical Solutions, Erlangen, Germany) using a 64-channel RF head coil. For the purposes of this study, only T1-weighted acquisitions were included. Table 1 provides a summary of the acquisition protocol parameters.

**Table 1.** A summary of the included T1-weighted anatomical MRI acquisition parameters included in the dataset and their respective counts.

| Scan Description | TE (ms) | TR (ms) | TI (ms) | Flip Angle (°) | X (mm) | Y (mm) | Z (mm) | Count |
| --- | --- | --- | --- | --- | --- | --- | --- | --- |
| Anatomy_iso1mm | 2.59 | 1750 | 900 | 8 | 1 | 1 | 1 | 1 |
|  | 2.61 | 1750 | 900 | 8 | 1 | 1 | 1 | 96 |
| MPRAGE_EnhancedContrast | 2.45 | 2530 | 1100 | 7 | 1 | 1 | 1 | 17 |
|  | 2.88 | 2530 | 1100 | 7 | 1 | 1 | 1 | 215 |
|  | 2.99 | 2530 | 1100 | 7 | 1 | 1 | 1 | 172 |
| MPRAGE_EnhancedContrast_MPR_Cor | 2.99 | 2530 | 1100 | 7 | 1 | 1 | 1 | 10 |
| MPRAGE_EnhancedContrast_MPR_Sag | 2.99 | 2530 | 1100 | 7 | 1 | 1 | 1 | 10 |
| MPRAGE_TFL | 2.9 | 2530 | 1100 | 7 | 1 | 1 | 1 | 18 |
| MPRAGE_iso0.6mm | 2.84 | 1750 | 900 | 8 | 0.597 | 0.597 | 0.6 | 100 |
| MPRAGE_iso1mm | 2.45 | 1750 | 900 | 8 | 1 | 1 | 1 | 16 |
|  | 2.59 | 1750 | 900 | 8 | 1 | 1 | 1 | 37 |
|  | 2.61 | 1750 | 900 | 8 | 1 | 1 | 1 | 165 |
| T1w_Anatomy_RL | 2.78 | 2400 | 1000 | 8 | 0.898 | 0.898 | 0.9 | 44 |
| T1w_MPRAGE_HCP | 2.14 | 2400 | 1000 | 8 | 1 | 1 | 1 | 10 |
| T1w_MPRAGE_RL | 2.77 | 2400 | 1000 | 8 | 1 | 1 | 1 | 6 |
|  | 2.78 | 2400 | 1000 | 8 | 0.875 | 0.875 | 0.9 | 162 |
|  | 2.78 | 2400 | 1000 | 8 | 0.898 | 0.898 | 0.9 | 2304 |
|  | 2.78 | 2400 | 1000 | 8 | 0.902 | 0.902 | 0.9 | 526 |
|  | 2.78 | 2400 | 1000 | 8 | 0.91 | 0.91 | 0.9 | 40 |
| t1_mp2rage_sag_p3_iso_INV1 | 3.52 | 4000 | 716 | 5 | 1 | 1 | 1 | 50 |
| t1_mp2rage_sag_p3_iso_INV2 | 3.52 | 4000 | 2180 | 5 | 1 | 1 | 1 | 50 |

### MRI Preprocessing

All scans were preprocessed using the FMRIB Software Library (FSL) v6.0 (see https://fsl.fmrib.ox.ac.uk/fsl/fslwiki/FSL). Brain extraction was applied using the BET script, with the configuration set to ‘robust’ (for more information, see https://fsl.fmrib.ox.ac.uk/fsl/fslwiki/BET/UserGuide). Following brain extraction, linear registration was applied with FLIRT using the 2 mm MNI152 template as the reference and the default affine registration mode with 12 degrees of freedom (for more information, see https://fsl.fmrib.ox.ac.uk/fsl/fslwiki/FLIRT/UserGuide). These two steps resulted in a dataset of 91?109?91 uniformly aligned volumes.

### Modeling

#### Architecture

The Keras-Tuner library was used for training and tuning the employed CNN architecture. The HyperXception (https://keras.io/api/keras_tuner/hypermodels/hyper_xception/) model was adapted from its existing implementation for multicategory classification to binary classification for the prediction of sex, and regression for the prediction of age and personality traits. The adapted version is openly accessible on GitHub at https://tinyurl.com/m7d2vr8e. This HyperModel subclass inherits from the original implementation, but includes several notable differences:

1. The original “top” layer, i.e., the output layer, which employs a Softmax activation for the prediction of a category, was dropped, and replaced with either a sigmoid activation (for sex classification) or a linear activation (for age and personality traits).
2. Before the addition of the new output layer, a vector of 7 elements representing scan acquisition parameters was concatenated to the CNN’s output. These included elements were: TR, TE, TI, flip angle, and resolution in the three axes. All elements were divided by constants in the required scale to produce values between 0 and 1.
3. The hyperparameter used to adjust the Adam optimization algorithm’s learning rate was slightly modified to check for a larger range of values (for more information about Adam, see Tuning, as well as https://keras.io/api/optimizers/adam/).
4. The loss function was changed from categorical cross-entropy to binary cross-entropy (BCE) for sex classification, mean absolute error (MAE) for age estimation, or mean squared error (MSE) for personality traits (averaged over the five outputs). The decision between MAE and MSE was made based on MAE’s advantage in interpretability versus MSE’s robustness to outliers (Willmott & Matsuura, 2005).

#### Tuning

The Keras-Tuner library’s Hyperband model (Li et al., 2018) was used for hyperparameter tuning (for more information, see https://keras.io/api/keras_tuner/tuners/hyperband/) with the tuner’s objective set to the validation loss (for more information on the chosen loss functions, see Architecture). The Adam optimizer (Kingma & Ba, 2017) was used in the training process of each selected hyperparameter configuration. Adam is a popular extension for the classic method of stochastic gradient descent (SGD), and has been shown to perform exceptionally well in a large variety of tasks (Llugsi et al., 2021; Ruder, 2017). Table 2 offers a complete overview of the evaluated hyperparameter values.

**Table 2.** Summary of the hyperparameter values which were evaluated during the Xception architecture’s tuning process.

| Parameter |  | Values |  |  |  |  |  |
| --- | --- | --- | --- | --- | --- | --- | --- |
| Activation |  | ReLU |  |  | SELU |  |  |
| Initial convolution | Number of filters | 32 |  | 64 |  | 128 |  |
|  | Kernel size | 3 |  |  | 5 |  |  |
| Residual blocks | Number of blocks | min: 2, max: 8, step: 1 |  |  |  |  |  |
|  | Number of filters | min: 128, max: 268, step: 128 |  |  |  |  |  |
| Pooling |  | average |  | flatten |  | max |  |
| Learning rate |  | 1e-2 | 1e-3 | 5e-4 | 1e-4 | 5e-5 | 1e-5 |

For sex and age, the validation loss was calculated for 395 of the 4,049 scans included in this study (∼9.75%), and no test set was used. The omission of the test set was decided upon with the intention of maximizing the training dataset and increasing the potential for selecting an architecture which will prove beneficial for its ultimate purpose, which will be its application for TL. For the estimation of personality traits, 200 of the 1,253 scans (belonging to 100 of the 415 total participants) were used as a validation set, and 155 scans (from 78 participants) were kept aside as a test set and not used in the training procedure in any way. All validation and test set scans were taken from subjects not included in the training sets.

As the tuning works in exploratory “brackets”, in which a subset of models is trained and evaluated, and then a smaller subset is selected for the consecutive bracket, several parameters were customized to increase the reliability of the results; the reduction factor between brackets was set to 3, each trial was configured to be executed twice, and the hyperband search itself was iterated twice. The duplication of each trial’s execution and hyperband iterations reduces the effect of randomly better or worse weights at initialization. All training set files were shuffled before each epoch to increase the robustness of the learning process.

### 5.2.5. Transfer Learning

For the application of TL, each model was loaded, and all existing layers were frozen (i.e., their weights were configured so that they would not be updated during training). The last layer, which was the prediction layer for the source task, was removed and replaced with a new prediction layer based on the tested target; a dense layer with a single unit using a sigmoid activation function for sex classification and a linear activation for age estimation, or five units with a linear activation function for personality traits. A dropout rate of 0.5 was introduced to reduce overfitting, and the Adam optimizer was used with a *1e*^*-5*^ learning rate and the same loss functions used in the original training procedures (see the Architecture section). The fitting routine was executed with an early stopping callback configured to wait for twenty epochs before termination and restoration of the best weights. After the model converged for the new prediction task, another “fine-tuning” iteration was made; all layers were unfrozen, the optimizer’s learning rate reduced to *5e*^*-7*^, and fitting restarted with the same objective and early stopping rule. See figure 2 for a general overview of the transfer learning procedure.

**Figure 2.**
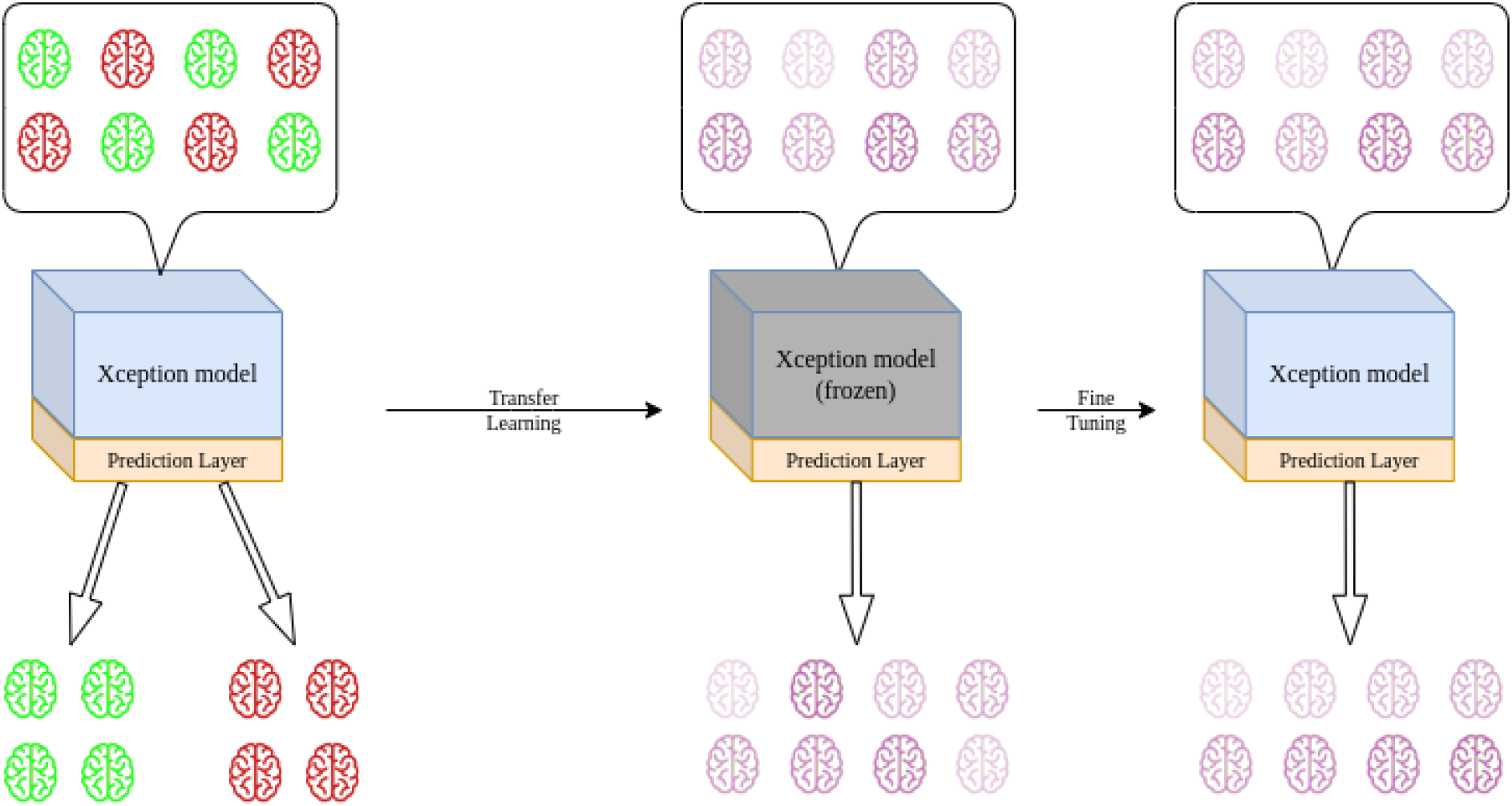
An illustration of the transfer learning (TL) procedure. This illustration depicts transfer from binary classification to regression; however, the same principle applies between any types of prediction tasks.

## 5.3. Results

### 5.3.1. Xception Tuning

Final results from the iterative hyperparameter tuning conducted by Keras-Tuner returned three highly distinct architectures. Table 3 summarizes the results from this procedure, as well as the calculated metrics from the included datasets. All selected Xception architectures displayed a significant degree of overfitting; sex classification accuracy over the training set scored 0.91, whereas the validation set achieved 0.83. Age prediction MAE over the training set was 1.523 years (R^2^=0.97) versus 4.348 years (R^2^=0.612) on the validation set. Finally, while the prediction of personality traits achieved slightly better average MSE on the validation set than on the training set, this was the result of the decreased variation in the sample. R^2^ values were slightly negative for both the validation and the test sets, indicating a “naive” model constantly predicting the average trait scores would have obtained a comparable result.

**Table 3.**
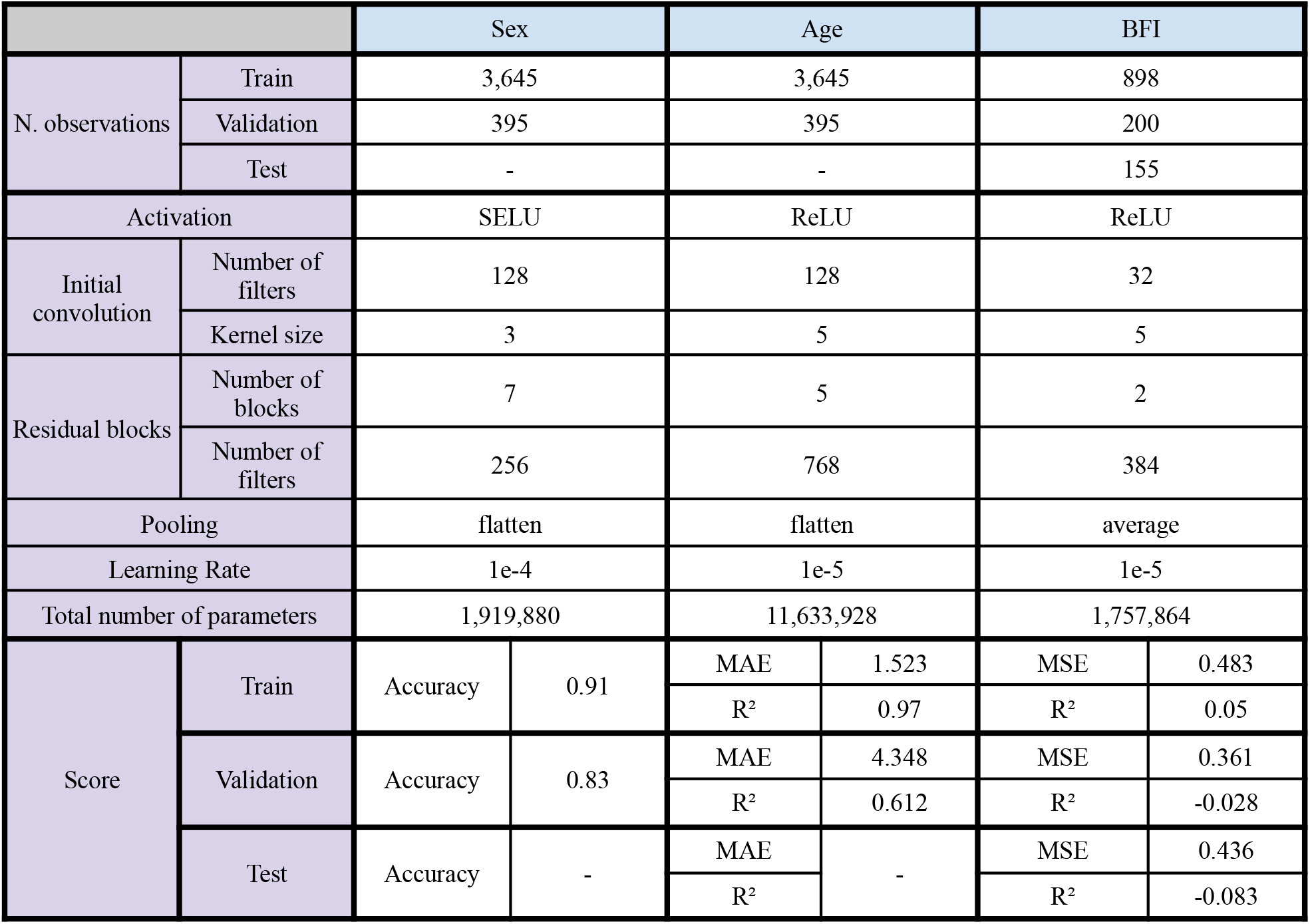
Hyperparameter tuning results. The top section details the number of observations included in each dataset. The middle section provides a summary of the selected hyperparameter values. The bottom section includes the calculated evaluation metrics for each dataset.

It is interesting to note that generally, very slow learning rates were found to be the most effective. Tuning for age estimation resulted in the largest network (i.e., the highest number of trainable parameters), over six times the size of the chosen network for sex classification. In addition, both the selected network architectures for sex and age used flattening as their pooling method preceding the “top” prediction layer. This is somewhat surprising, as flattening considerably increases the number of network parameters, and is significantly less frequently used in practical applications.

### 5.3.2. Transfer Learning

The trained models sex and age have been retrained and evaluated both for the complementary domain, as well as for the prediction of personality traits (see figure 2 for an overview of the TL retraining procedure). Our results indicate that TL applied between sex and age achieved comparable results, despite the considerable difference in network architecture. The larger, age prediction network even improved upon the selected sex classification architecture, but also resulted in more overfitting (training set accuracies of 0.91 vs. 1 and validation set accuracies of 0.83 vs. 0.86).

The prediction of BFI personality traits did not show any improvement over the non-TL training procedure. Both trained sex and age prediction network architectures returned negative R^2^ values, indicating that the prediction MSE would be lower if the model simply returned the average BFI scores for each trait. Figure 3 and table 4 provide a summary of the transfer learning training procedure and the results.

**Table 4.** TL performance summary. Loss metrics are emphasized in bold lettering.

| Source | Target | Sex |  |  |  | Age |  |  |  | BFI |  |  |  |
| --- | --- | --- | --- | --- | --- | --- | --- | --- | --- | --- | --- | --- | --- |
|  | Metric | **BCE** |  | Accuracy |  | **MAE** |  | R2 |  | **MSE** |  | R2 |  |
|  | Evaluation Set | Train | Validation | Train | Validation | Train | Validation | Train | Validation | Train | Validation | Train | Validation |
| Sex |  |  |  |  |  | 3.02 | 4.46 | 0.73 | 0.58 | 0.04 | 0.44 | 0.89 | -0.27 |
| Age |  | 0.02 | 0.39 | 1 | 0.86 |  |  |  |  | 0.46 | 0.56 | 0.32 | -0.64 |

**Figure 3.**
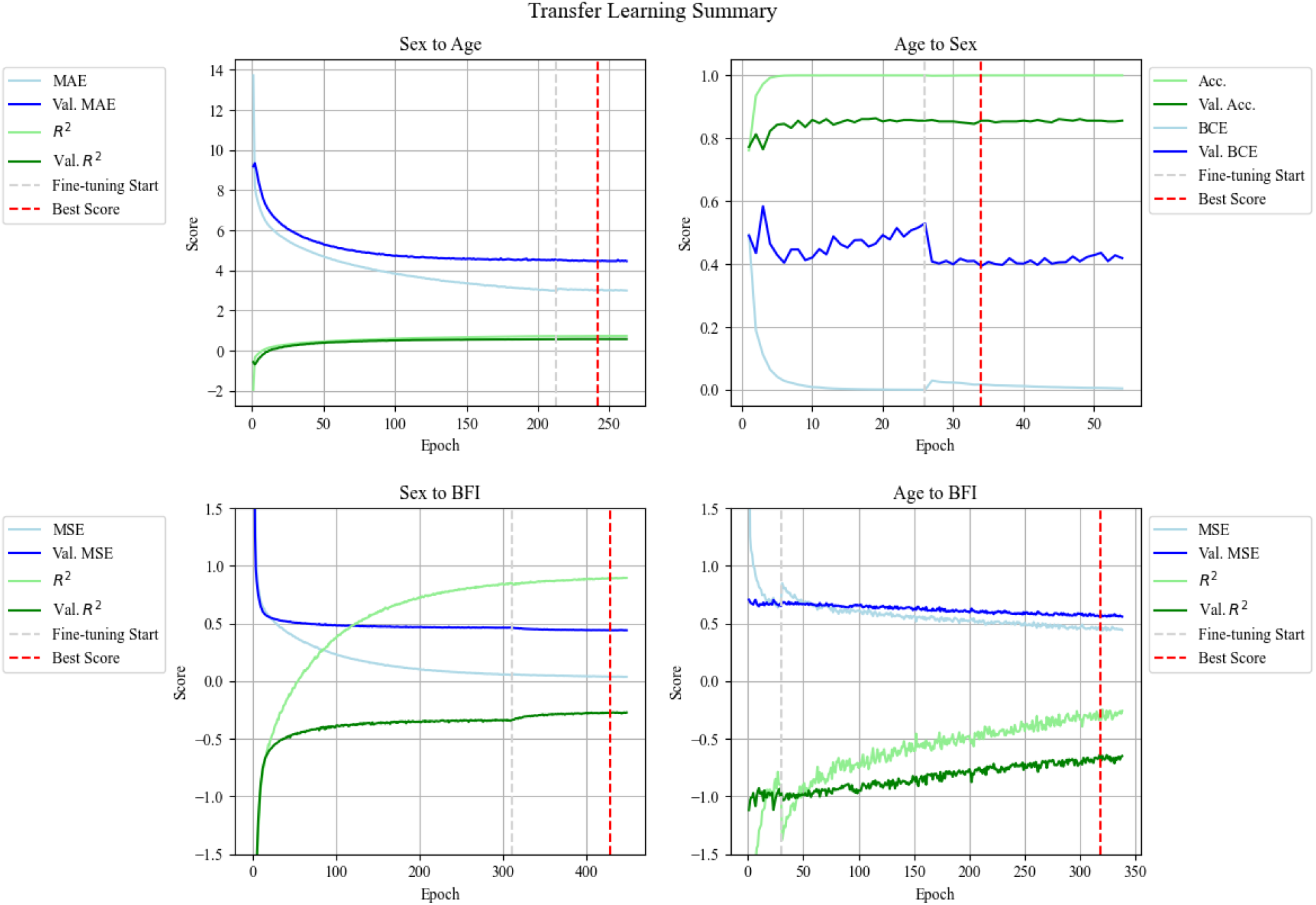
A summary of all TL training procedures. Blue lines indicate training and validation loss (MAE for age, BCE for sex, and MSE for BFI scores). Green lines indicate an additional metric (accuracy in sex classification, R^2^ in age and BFI scores prediction).

## 5.4. Discussion

The field of neuroimaging, together with many other scientific domains, is facing a dramatic transition in terms of the scale of available data and the computational resources available to leverage them. This transition holds the promise of unprecedented statistical power to detect and study the finest relationships between measurable brain features and various aspects of human function and cognition. However, together with the technological advancements that enable these developments, new challenges arise.

In this study, we tested the performance of an adapted version of an automatic Xception CNN hyperparameter tuning algorithm, HyperXception, in predicting sex and age using a relatively large dataset of 4,049 T1-weighted anatomical MRI scans from 1,594 participants. Furthermore, we tested the performance of the algorithm in predicting personality traits from a smaller sample (1,098 scans from 337 participants, and an additional test set of 155 from 78 participants), as represented by BFI scores. Finally, we compared the results of the selected network hyperparameter configurations for the prediction of age and sex with the results attained by implementing a TL workflow over its respective counterpart, as well as over BFI scores.

### 5.4.1. Interpretation

TL applied between sex and age resulted in comparable prediction performance metrics, despite significant differences in the respective networks’ architecture. These results indicate that both networks were successful in the extraction of informative features from the provided T1-weighted sMRI dataset. It is possible that performance might still be improved using different architectures, various data augmentation techniques, different preprocessing, etc. However, the focus of this article is not the optimization of sex or age prediction, rather, the evaluation of learning transferability between domains when applied following a uniform preprocessing procedure. In this respect, the results demonstrate the potential of CNNs to become a useful tool for sharing informative encodings for large datasets.

The failure in predicting BFI scores may be attributed to several possible reasons, with the most likely, in our estimation, being either that these effects require much more data, or that personality traits simply cannot be estimated from T1-weighted sMRI. Likewise, while BFI scores are considered relatively reliable, it is possible that a qualitative self-reporting questionnaire is too noisy for a model to effectively learn generalizable weight values over any feasibly large sample.

### 5.4.2. Limitations and Future Work

Although the core principle evaluated in this study is applicable across architectures and domains, we note that this procedure should still be applied to more CNNs and different target variables to develop a finer understanding of the method’s limitations. The Keras-Tuner library already provides a HyperResNet model (see https://keras.io/api/keras_tuner/hypermodels/hyper_resnet/) which may be used, and more architectures are likely to be added in the near future. In addition, it would be interesting to evaluate the results attained by 3D CNN architectures leveraging several modalities at once (e.g., an extended sMRI network utilizing T1-weighted, T2-weighted, and FLAIR acquisitions simultaneously).

A major limitation of this study is the sample size and the omission of a test set of the sex and age samples. It would be beneficial to apply the same methodology to a sample size at the scale of the UK BioBank (over 50,000 participants) and re-evaluate the results. Another aspect which requires further investigation is the influence of the acquisition site and parameters. The dataset used for the purposes of this exploration was collected from a single scanner and biased towards a particular scanning protocol (2,304 of the 4,049 scans). Integration of different datasets representing a diverse sample of sites, machines, and acquisition protocols, would significantly improve the reliability of the results. All the aforementioned variables have a demonstrated contribution to variation in aggregated data (J. Chen et al., 2014; Han et al., 2006; Jovicich et al., 2006; Kostro et al., 2014; Streitbürger et al., 2014). Given a large enough sample, it would be appealing to include these parameters with the encoded volumetric data and assess their importance in a variety of prediction tasks.

## References

Akkus, Z., Galimzianova, A., Hoogi, A., Rubin, D. L., & Erickson, B. J. (2017). Deep Learning for Brain MRI Segmentation: State of the Art and Future Directions. Journal of Digital Imaging, 30(4), 449–459. https://doi.org/10.1007/s10278-017-9983-4

Andrade, N., Faria, F., & Cappabianco, F. (2018). A Practical Review on Medical Image Registration: From Rigid to Deep Learning Based Approaches (p. 470). https://doi.org/10.1109/SIBGRAPI.2018.00066

Badrinarayanan, V., Kendall, A., & Cipolla, R. (2017). SegNet: A Deep Convolutional Encoder-Decoder Architecture for Image Segmentation. IEEE Transactions on Pattern Analysis and Machine Intelligence, 39(12), 2481–2495. https://doi.org/10.1109/TPAMI.2016.2644615

Bycroft, C., Freeman, C., Petkova, D., Band, G., Elliott, L. T., Sharp, K., Motyer, A., Vukcevic, D., Delaneau, O., O’Connell, J., Cortes, A., Welsh, S., Young, A., Effingham, M., McVean, G., Leslie, S., Allen, N., Donnelly, P., & Marchini, J. (2018). The UK Biobank resource with deep phenotyping and genomic data. Nature, 562(7726), Article 7726. https://doi.org/10.1038/s41586-018-0579-z

Chen, J., Liu, J., Calhoun, V. D., Arias-Vasquez, A., Zwiers, M. P., Gupta, C. N., Franke, B., & Turner, J. A. (2014). Exploration of scanning effects in multi-site structural MRI studies. Journal of Neuroscience Methods, 230, 37–50. https://doi.org/10.1016/j.jneumeth.2014.04.023

Chen, S., Ma, K., & Zheng, Y. (2019). Med3D: Transfer Learning for 3D Medical Image Analysis (arXiv:1904.00625). arXiv. http://arxiv.org/abs/1904.00625

Chen, X., Diaz-Pinto, A., Ravikumar, N., & Frangi, A. F. (2021). Deep learning in medical image registration. Progress in Biomedical Engineering, 3(1), 012003. https://doi.org/10.1088/2516-1091/abd37c

Chollet, F. (2017). Xception: Deep Learning with Depthwise Separable Convolutions. 2017 IEEE Conference on Computer Vision and Pattern Recognition (CVPR), 1800–1807. https://doi.org/10.1109/CVPR.2017.195

Dalca, A. V., Yu, E., Golland, P., Fischl, B., Sabuncu, M. R., & Iglesias, J. E. (2019). Unsupervised Deep Learning for Bayesian Brain MRI Segmentation. Medical Image Computing and Computer-Assisted Intervention : MICCAI … International Conference on Medical Image Computing and Computer-Assisted Intervention, 11766, 356–365. https://doi.org/10.1007/978-3-030-32248-9_40

de Vos, B. D., Berendsen, F. F., Viergever, M. A., Sokooti, H., Staring, M., & Isgum, I. (2019). A Deep Learning Framework for Unsupervised Affine and Deformable Image Registration. Medical Image Analysis, 52, 128–143. https://doi.org/10.1016/j.media.2018.11.010

Di Martino, A., O’Connor, D., Chen, B., Alaerts, K., Anderson, J. S., Assaf, M., Balsters, J. H., Baxter, L., Beggiato, A., Bernaerts, S., Blanken, L. M. E., Bookheimer, S. Y., Braden, B. B., Byrge, L., Castellanos, F. X., Dapretto, M., Delorme, R., Fair, D. A., Fishman, I., … Milham, M. P. (2017). Enhancing studies of the connectome in autism using the autism brain imaging data exchange II. Scientific Data, 4, 170010. https://doi.org/10.1038/sdata.2017.10

Di Martino, A., Yan, C.-G., Li, Q., Denio, E., Castellanos, F. X., Alaerts, K., Anderson, J. S., Assaf, M., Bookheimer, S. Y., Dapretto, M., Deen, B., Delmonte, S., Dinstein, I., Ertl-Wagner, B., Fair, D. A., Gallagher, L., Kennedy, D. P., Keown, C. L., Keysers, C., … Milham, M. P. (2014). The autism brain imaging data exchange: Towards a large-scale evaluation of the intrinsic brain architecture in autism. Molecular Psychiatry, 19(6), Article 6. https://doi.org/10.1038/mp.2013.78

Frei, J., Frei-Stuber, L., & Kramer, F. (2022). GERNERMED++: Transfer Learning in German Medical NLP arXiv:2206.14504). arXiv. http://arxiv.org/abs/2206.14504

Girshick, R. (2015). Fast R-CNN (arXiv:1504.08083). arXiv. https://doi.org/10.48550/arXiv.1504.08083

Godasu, R., Zeng, D., & Sutrave, K. (2020). Transfer Learning in Medical Image Classification: Challenges and Opportunities.

Gore, D. V., & Deshpande, V. (2020). Comparative Study of various techniques using Deep Learning for Brain Tumor Detection. 2020 International Conference for Emerging Technology (INCET), 1–4. https://doi.org/10.1109/INCET49848.2020.9154030

Han, X., Jovicich, J., Salat, D., van der Kouwe, A., Quinn, B., Czanner, S., Busa, E., Pacheco, J., Albert, M., Killiany, R., Maguire, P., Rosas, D., Makris, N., Dale, A., Dickerson, B., & Fischl, B. (2006). Reliability of MRI-derived measurements of human cerebral cortical thickness: The effects of field strength, scanner upgrade and manufacturer. NeuroImage, 32(1), 180–194. https://doi.org/10.1016/j.neuroimage.2006.02.051

He, K., Zhang, X., Ren, S., & Sun, J. (2016). Deep Residual Learning for Image Recognition. 2016 IEEE Conference on Computer Vision and Pattern Recognition (CVPR), 770–778. https://doi.org/10.1109/CVPR.2016.90

Jovicich, J., Czanner, S., Greve, D., Haley, E., van der Kouwe, A., Gollub, R., Kennedy, D., Schmitt, F., Brown, G., MacFall, J., Fischl, B., & Dale, A. (2006). Reliability in multi-site structural MRI studies: Effects of gradient non-linearity correction on phantom and human data. NeuroImage, 30(2), 436–443. https://doi.org/DOI:10.1016/j.neuroimage.2005.09.046

Kingma, D. P., & Ba, J. (2017). Adam: A Method for Stochastic Optimization (arXiv:1412.6980). arXiv. http://arxiv.org/abs/1412.6980

Kora, P., Ooi, C. P., Faust, O., Raghavendra, U., Gudigar, A., Chan, W. Y., Meenakshi, K., Swaraja, K., Plawiak, P., & Rajendra Acharya, U. (2022). Transfer learning techniques for medical image analysis: A review. Biocybernetics and Biomedical Engineering, 42(1), 79–107. https://doi.org/10.1016/j.bbe.2021.11.004

Kostro, D., Abdulkadir, A., Durr, A., Roos, R., Leavitt, B. R., Johnson, H., Cash, D., Tabrizi, S. J., Scahill, R. I., Ronneberger, O., & Klöppel, S. (2014). Correction of inter-scanner and within-subject variance in structural MRI based automated diagnosing. NeuroImage, 98, 405–415. https://doi.org/10.1016/j.neuroimage.2014.04.057

Li, L., Jamieson, K., DeSalvo, G., Rostamizadeh, A., & Talwalkar, A. (2018). Hyperband: A Novel Bandit-Based Approach to Hyperparameter Optimization. Journal of Machine Learning Research, 18, 1–52.

Liem, F., Varoquaux, G., Kynast, J., Beyer, F., Kharabian Masouleh, S., Huntenburg, J. M., Lampe, L., Rahim, M., Abraham, A., Craddock, R. C., Riedel-Heller, S., Luck, T., Loeffler, M., Schroeter, M. L., Witte, A. V., Villringer, A., & Margulies, D. S. (2017). Predicting brain-age from multimodal imaging data captures cognitive impairment. NeuroImage, 148, 179–188. https://doi.org/10.1016/j.neuroimage.2016.11.005

Liu, M., Zhang, J., Lian, C., & Shen, D. (2020). Weakly-supervised Deep Learning for Brain Disease Prognosis using MRI and Incomplete Clinical Scores. IEEE Transactions on Cybernetics, 50(7), 3381–3392. https://doi.org/10.1109/TCYB.2019.2904186

Llugsi, R., Yacoubi, S. E., Fontaine, A., & Lupera, P. (2021). Comparison between Adam, AdaMax and Adam W optimizers to implement a Weather Forecast based on Neural Networks for the Andean city of Quito. 2021 IEEE Fifth Ecuador Technical Chapters Meeting (ETCM), 1–6. https://doi.org/10.1109/ETCM53643.2021.9590681

Lundervold, A. S., & Lundervold, A. (2019). An overview of deep learning in medical imaging focusing on MRI. Zeitschrift Für Medizinische Physik, 29(2), 102–127. https://doi.org/10.1016/j.zemedi.2018.11.002

Nazir, M., Shakil, S., & Khurshid, K. (2021). Role of deep learning in brain tumor detection and classification (2015 to 2020): A review. Computerized Medical Imaging and Graphics, 91, 101940. https://doi.org/10.1016/j.compmedimag.2021.101940

Noor, M. B. T., Zenia, N. Z., Kaiser, M. S., Mahmud, M., & Al Mamun, S. (2019). Detecting Neurodegenerative Disease from MRI: A Brief Review on a Deep Learning Perspective. In P. Liang, V. Goel, & C. Shan (Eds.), Brain Informatics (pp. 115–125). Springer International Publishing. https://doi.org/10.1007/978-3-030-37078-7_12

Peng, Y., Yan, S., & Lu, Z. (2019). Transfer Learning in Biomedical Natural Language Processing: An Evaluation of BERT and ELMo on Ten Benchmarking Datasets (arXiv:1906.05474). arXiv. http://arxiv.org/abs/1906.05474

Ruder, S. (2017). An overview of gradient descent optimization algorithms (arXiv:1609.04747). arXiv. http://arxiv.org/abs/1609.04747

Saratxaga, C. L., Moya, I., Picón, A., Acosta, M., Moreno-Fernandez-de-Leceta, A., Garrote, E., & Bereciartua-Perez, A. (2021). MRI Deep Learning-Based Solution for Alzheimer’s Disease Prediction. Journal of Personalized Medicine, 11(9), Article 9. https://doi.org/10.3390/jpm11090902

Shao, W., Banh, L., Kunder, C. A., Fan, R. E., Soerensen, S. J. C., Wang, J. B., Teslovich, N. C., Madhuripan, N., Jawahar, A., Ghanouni, P., Brooks, J. D., Sonn, G. A., & Rusu, M. (2021). ProsRegNet: A deep learning framework for registration of MRI and histopathology images of the prostate. Medical Image Analysis, 68, 101919. https://doi.org/10.1016/j.media.2020.101919

Siar, M., & Teshnehlab, M. (2019). Brain Tumor Detection Using Deep Neural Network and Machine Learning Algorithm (p. 368). https://doi.org/10.1109/ICCKE48569.2019.8964846

Streitbürger, D.-P., Pampel, A., Krueger, G., Lepsien, J., Schroeter, M. L., Mueller, K., & Möller, H. E. (2014). Impact of image acquisition on voxel-based-morphometry investigations of age-related structural brain changes. NeuroImage, 87, 170–182. https://doi.org/10.1016/j.neuroimage.2013.10.051

Torrey, L., & Shavlik, J. (2009). Transfer Learning. In Handbook of research on machine learning applications and trends: Algorithms, methods, and techniques (Vol. 1–2, pp. 242–264). IGI global. https://ftp.cs.wisc.edu/machine-learning/shavlik-group/torrey.handbook09.pdf

Van Essen, D. C., Smith, S. M., Barch, D. M., Behrens, T. E. J., Yacoub, E., & Ugurbil, K. (2013). The WU-Minn Human Connectome Project: An Overview. NeuroImage, 80, 62–79. https://doi.org/10.1016/j.neuroimage.2013.05.041

Van Essen, D. C., Ugurbil, K., Auerbach, E., Barch, D., Behrens, T. E. J., Bucholz, R., Chang, A., Chen, L., Corbetta, M., Curtiss, S. W., Della Penna, S., Feinberg, D., Glasser, M. F., Harel, N., Heath, A. C., Larson-Prior, L., Marcus, D., Michalareas, G., Moeller, S., … Yacoub, E. (2012). The Human Connectome Project: A data acquisition perspective. NeuroImage, 62(4), 2222–2231. https://doi.org/10.1016/j.neuroimage.2012.02.018

Willmott, C., & Matsuura, K. (2005). Advantages of the mean absolute error (MAE) over the root mean square error (RMSE) in assessing average model performance. Climate Research, 30, 79–82. https://doi.org/10.3354/cr030079

Wu, J. (2017). Introduction to Convolutional Neural Networks. National Key Lab for Novel Software Technology, 5(23), 495.

Xiao, H., Teng, X., Liu, C., Li, T., Ren, G., Yang, R., Shen, D., & Cai, J. (2021). A review of deep learning-based three-dimensional medical image registration methods. Quantitative Imaging in Medicine and Surgery, 11(12), 4895–4916. https://doi.org/10.21037/qims-21-175

Zhang, K., Zuo, W., Chen, Y., Meng, D., & Zhang, L. (2017). Beyond a Gaussian Denoiser: Residual Learning of Deep CNN for Image Denoising. IEEE Transactions on Image Processing, 26(7), 3142–3155. https://doi.org/10.1109/TIP.2017.2662206

